# Diversity in transcriptomics without cell types

**DOI:** 10.64898/2026.07.13.735796

**Authors:** Leyi Jiang, Katherine Benjamin, Jesse V Veenvliet, Emily Roff, Heather A Harrington

## Abstract

Downstream analysis in single-cell and spatial transcriptomics is highly dependent on a sequence of upstream modeling choices. The non-canonicity of these choices presents challenges for reproducibility. In particular, measures of cellular heterogeneity and diversity do not solely reflect biological variation, but are also sensitive to parameter settings. A diversity measure that is robust to modeling choices, such as clustering resolution, is therefore desirable to improve reproducibility and interpretability. Here, we introduce scDIV, a similarity-sensitive measure of cellular diversity inspired by mathematical ideas in ecological science, which is robust to graph-based clustering parameters and remains applicable even in the absence of cell-type clusters. We use scDIV to quantitatively track the progress of tissue differentiation in both single-cell and spatial mouse development datasets and to evaluate different engineered stem-cell-based embryo models. In contrast to traditional entropy-based methods, such as the Hill number, used to quantify biodiversity, scDIV remains robust to clustering.

## Introduction

Biological systems are remarkably robust [1, 2]. Understanding this robustness from the viewpoint of gene regulation has motivated the rapid generation of single-cell and spatial transcriptomic atlases across tissues and organisms [3–5]. However, the robustness of biological systems stands in stark contrast to current data analysis pipelines, which remain fragile in the face of increasing data complexity and sensitive to analytical and parameter choices [6]. To handle this ever-increasing complexity, new mathematical frameworks are required that can naturally and robustly capture the underlying geometry and topology of gene expression space. Rather than treating cell populations as discrete, isolated clusters, such a framework should ideally incorporate similarity and magnitude [7, 8] across multiple resolutions [9] to achieve a multiscale representation of biological systems.

Assigning cell types to transcriptomics data sets is a notoriously fragile and ill-defined task. This fragility is primarily driven by the upstream graph-based-clustering process, which relies on a series of sensitive parameter settings and analytical choices, including the choice of initial dimensionality reduction, the number of principal components to retain, and the clustering-resolution parameter. Even if a preprocessing pipeline is fixed, it has been shown that the choice of software library and its specific version can result in significant differences in downstream cell-type assignments [6]. This non-canonicity presents a problem: how do we design methods that are robust to the choice of cell-type clustering?

Cell-type diversity and heterogeneity measures are known to correlate with developmental processes, tissue function, regenerative capacity [10], and disease progression [11, 12]. During embryogenesis, initially similar cells give rise to increasingly complex tissues with distinct cell types and spatial arrangements. Similar principles govern organ development, tissue homeostasis, and regeneration, whereas disruptions of these processes can result in pathological states such as cancer and fibrosis. Quantifying and comparing how cellular diversity and heterogeneity unfolds in development, homeostasis and regeneration – and in health and disease – is pivotal for understanding the emergence, maintenance, and breakdown of multicellular organization. To this end, entropy-based methods have been proposed [13, 14]. For instance, Shannon entropy has successfully detected critical state transitions during erythroid progenitor differentiation [15] and served as a prognostic indicator in breast cancer immunology [11, 12]. More recently, Karagiannis et al. [16] adapted these concepts to single-cell transcriptomics. However, all of these entropy-based methods take only the cell-type distribution as input, and are therefore highly sensitive to reassignments and subclustering.

Here we propose scDIV, a similarity-sensitive framework for transcriptomic diversity inspired by the ecological diversity measures of Leinster and Cobbold [7]. The core idea in scDIV is the *Leinster-Cobbold (LC) diversity*, computed from both the relative abundances of the cell types and their pairwise similarities (Figure 1A). LC diversity, which is more robust to reclassification artifacts such as subclustering when compared to entropy-based measures like the Hill number (Figure 1B), gives a measure of the “effective number of cell types” in the sample (Figure 1C). Furthermore, in spatial data we can adapt LC diversity to reveal both the local and global diversity contributions of spatial subregions (Figure 1D).

**Figure 1.**
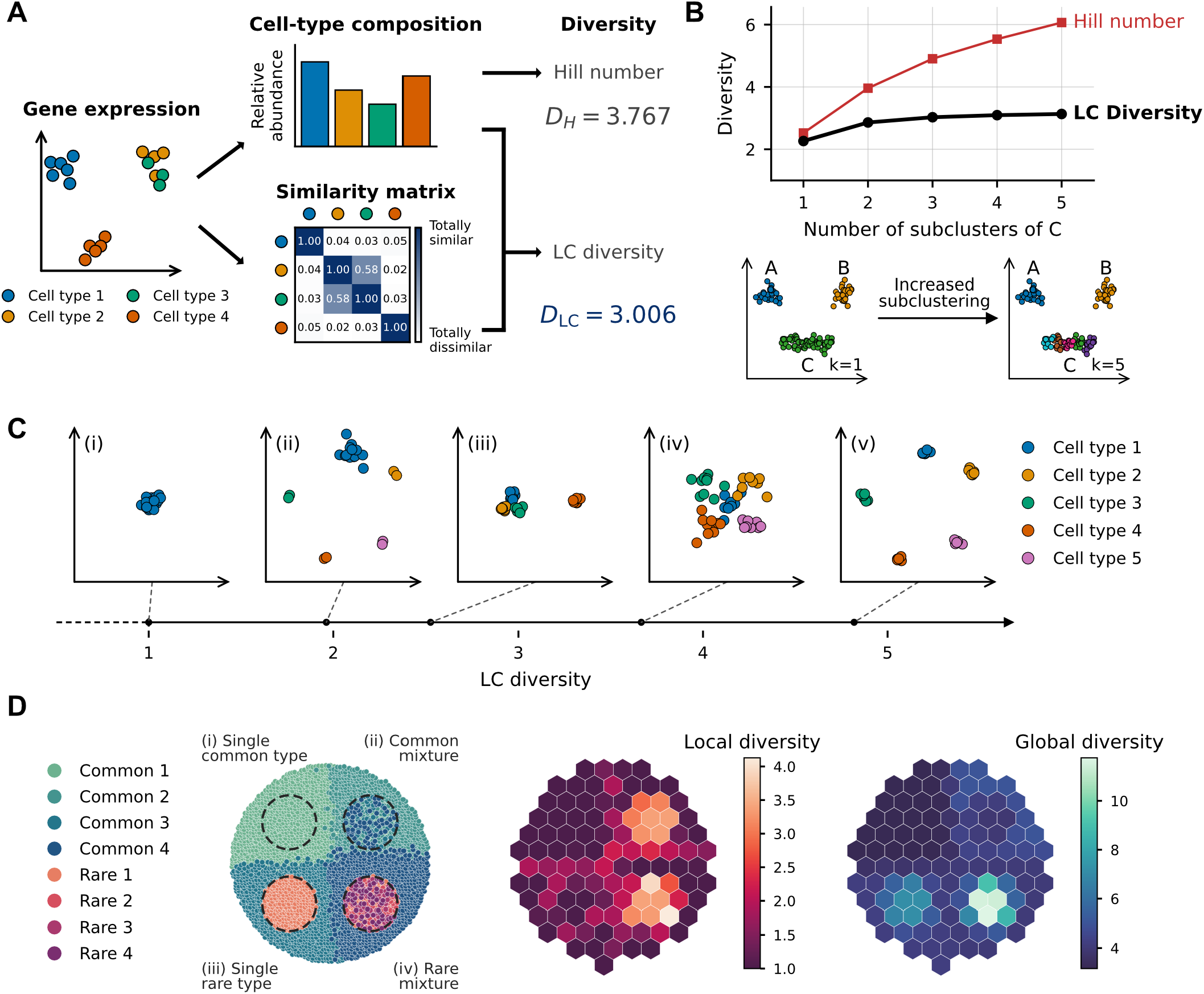
Method overview Low-dimensional schematics of single-cell data are used here for pedagogical purposes. (A) Traditional measures of diversity, such as the Hill number, only take into account the relative abundance of cell types. LC diversity also inputs the relative similarities of the cell types, providing robustness to different cell-type classifications. (B) LC diversity is robust to common choices in cell-type classification. Here the subcluster C is subdivided into increasingly many subclusters. Measuring diversity just from the cell-type composition, for example with Hill numbers, gives an overestimate of the true diversity of the underlying sample. LC diversity is robust, and remains stable under increasing subdivision. (C) LC diversity gives a measure of the *effective number of cell types* in a sample. (i) A single cell type has an LC diversity of 1. (ii) The addition of a handful of rare cells increases diversity a small amount, but LC diversity can see that the sample is dominated by a single cell type. (iii) Subdividing one cell type into several subtypes results in only a modest increase in the LC diversity score, here from ∼2 to ∼2.5. (iv) 5 cell types with high within-cell-type variance and poor separation have an LC diversity of less than 5. (v) 5 tight, well-separated clusters give an LC diversity of ∼5. (D) Local and global diversity measure of spatial data. Left: an example spatial data set with four regions of interest: (i) a region with a single globally common cell type; (ii) a region with a mixture of different globally common cell types; (iii) a region with a single globally rare cell type; (iv) a region with a mixture of different globally rare cell types. Middle: The local diversity map of the data set. The two mixture regions have high local diversities, whereas the two regions with only a single cell type have low local diversities. Additionally, the borders between different regions also exhibit higher local diversities because they represent mixtures of the tissue types in each region. Right: The global diversity map of the data set. The two rare regions have high global diversities, whereas the two regions with only common cell types have low global diversities.

We demonstrate the application of scDIV in a single-cell mouse development atlas, where the metric captures temporal diversification while remaining robust to clusterings. Using mouse spatial transcriptomics, we partition LC diversity into local and global components to map the emerging regional specialization of developing organs. Finally, in engineered stem-cell-based embryo models, the metric tracks transcriptomics shifts driven by extra-embryonic interactions. Together, these applications establish scDIV as a robust, pipeline-agnostic framework for quantifying cellular heterogeneity.

## Materials and Methods

### Similarity-sensitive diversity

We begin with an overview of the basic machinery of LC diversity [7], which is the central measure used by scDIV, before we show its application to transcriptomics data. A good general reference in the context of ecology is the textbook of Leinster [17].

A *community* (*S, p, Z*) *of size N* consists of the following data:

1. A list of *N members S* = [*S*_1_, …, *S*_*N*_].
2. A distribution *p* ∈ [0, 1]^*N*^ such that *p*_*i*_ measures the *relative abundance* of the member *S*_*i*_ in the community.
3. An *N ×N* symmetric matrix *Z* with entries *Z*_*ij*_ ∈ [0, 1] measuring the *similarity* between members *S*_*i*_ and *S*_*j*_ .

We require that the distribution satisfie 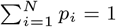. We will more precisely relate these notions to transcriptomics shortly, but for now a working intuition is to think of *S* as being the list of cell types assigned to some transcriptomics data.

The classical way of measuring the diversity of a community is to look only at the relative abundance distribution *p*. In single-cell transcriptomics, this is often done by simple visual inspection, but one can also quantify this intuition via the *Hill number* of the distribution [18]. For a distribution *p* ∈ [0, 1]^*N*^, its *Hill number* is

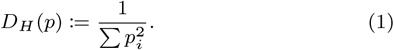

One can interpret the Hill number as describing the *effective number of members* of a community. Indeed, a community with only a single active member (*p*_*i*_ = 1 for some *i*) will have Hill number *D*_*H*_ (*p*) = 1. On the other extreme, a community with a perfectly even distribution of members (*p*_*i*_ = 1*/N* for all *i*) will have Hill number *D*_*H*_ (*p*) = *N*.

However, we argue that the similarity matrix *Z* carries essential information for quantifying diversity of a transcriptomics sample. In particular, it allows for a distinction to be made between two truly distinct cell types and two types which are merely subtypes of some supertype. The similarity-sensitive diversity measure of Leinster and Cobbold [7] is precisely what we need. For a community (*S, p, Z*), its *LC diversity* is

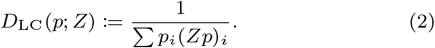

Now the contribution of each member *S*_*i*_ has been weighted by the value 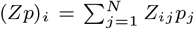, which measures the *expected similarity of i* to a uniformly selected member of the community.

The LC diversity can still be interpreted as the effective number of members in the community (see Figure 1C). However, in contrast to the Hill number, the similarity weighting gives it a robustness to re-labelings and, in particular, to repeated subtyping. In Figure 1B we can see how repeated subtyping heavily distorts the Hill number while the LC diversity remains stable at approximately 3 effective cell types. In fact, the LC diversity is a direct generalization of the Hill number; we have *D*_LC_(*p*; *I*) = *D*_*H*_ (*p*) for any *p*, so the Hill number can be viewed as the LC diversity given complete pairwise dissimilarity of community members.

For further discussion on LC diversity refer to Supplementary Information S1. This note covers the *sensitivity parameter*, which is fixed in this study but can be adjusted to control for sensitivity to rare cell types, and formal results on stability of LC diversity with respect to reclassification.

### Capturing diversity in transcriptomics

To compute the LC diversity of a single-cell transcriptomics data set, two pieces of information need to be input: an abundance distribution of cell types and a pairwise similarity matrix between cell types. Throughout this text, we will use terms ‘cell-type annotation,’ ‘assignment,’ and ‘clustering’ interchangeably for readability. However, a distinction must be made between the computational step of graph-based clustering and the subsequent biological step of naming the resulting clusters. While our diversity measures depend on the underlying cluster partition, they are agnostic to the biological labels of cell types.

The abundance distribution can be derived directly from the cell clusters generated by a standard single-cell analysis pipeline. An alternative made possible by the addition of similarity data is to operate in *singleton* mode, where each cell represents its own unique member within the population. The advantage of this approach is that the user need not worry about the sensitive upstream choices of modeling parameters, such as clustering resolution, of the cell-type annotation pipeline.

Once a community of cell types has been defined, a pairwise similarity matrix must be computed between them. In this paper, we compute similarity between two cell types *S*_*i*_ and *S*_*j*_ as a cosine similarity between their power-transformed average expression vectors. We first set a parameter *α* which controls for the relative importance of highly-expressed genes. Then, if the two cell types have average expression vectors *X*_*i*_ and *X*_*j*_ respectively, their similarity is

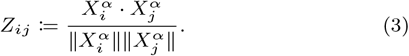

The parameter *α* allows us to control for the effect of any housekeeping genes which dominate expression but are relatively low variance. In this paper we set *α* = 0.5 mirroring the Bhattacharyya distance as previously studied [19].

### Diversity in spatial transcriptomics

For spatial transcriptomics, we are interested in the diversity of subregions of some larger spatial data set. Here we follow the diversity-partitioning framework of Reeve et al. [20]. There are two natural ways to assign these measurements, which we will call *local* and *global* diversity. In *local* diversity, we measure the diversity of the region as its own distinct unit. So, heterogeneous regions will have a high local diversity and homogeneous regions will have a low local diversity. In *global* diversity, we calculate the contribution of the region *to the spatial sample as a whole*. So, a region that contains unique cells relative to the wider cellular population will have a high global diversity, even if it has a low local diversity. See Supplementary Information S1.3.

Some spatial transcriptomics technologies, especially those with high resolution such as Stereo-seq [21] and Slide-seqV2 [22], display high numbers of dropped genes. To protect against distortion of the diversity statistic in these contexts, we apply a smoothing procedure by averaging each spatial unit with its nearest neighbors in PCA space. We then partition the slide into hexagonal regions and drop any regions with a low absolute cell count or a low density of cells.

### Implementation

We have implemented scDIV as a Python software package for similarity-sensitive transcriptomic diversity analysis. For single-cell data, `scdiv` exposes two modes: cell-type mode, and singleton mode. In cell-type mode, user-defined cell types are used as the fundamental members in the LC computation. In singleton mode, each individual cell is treated as a unique member with a uniform distribution over all members. For spatial transcriptomics, `scdiv` features a spatial module that automatically partitions tissue coordinates into square or hexagonal geometric regions and computes local and global diversity across these spatial regions.

### Data processing

We applied diversity computation to a collection of transcriptomics data sets, including mouse development single-cell atlas [23], Mouse Organogenesis Spatiotemporal Transcriptomic Atlas (MOSTA) [21] and a time-series scRNA-seq of engineered mouse stem-cell-based embryo models [24].

The mouse development data set [23] consists of 11 400 000 nuclei derived from 74 embryos, sampled at 6 h intervals across 43 timepoints spanning late gastrulation (embryonic day (E)8) to birth (postnatal day (P)0). The data has already been quality controlled according to Qiu et al. [23]. We further identified the 2000 most highly variable genes (HVGs) per embryonic stage using the Seurat `FindVariableFeatures` function and then retained only those genes identified as variable in more than three independent timepoints. This filtering resulted in a final set of 3754 HVGs, which were used to compute all diversity measures. In addition, we performed *de novo* clustering on the data set. First, expression counts were normalized, log-transformed, and projected onto the top 50 principal components. These components were used to construct a *k*-nearest-neighbor graph (*k* = 20). Finally, cells were partitioned via Leiden clustering with a resolution of 0.5 over 2 iterations.

MOSTA uses Stereo-seq to capture developmental dynamics at sub-cellular resolution across embryonic days E9.5 to E16.5 [21]. From this atlas, we obtained the processed data set at bin50 resolution (approximately 25 µm center-to-center bin distance) of a continuous developmental series of eight whole-embryo sagittal sections sampled at 1-day intervals spanning E9.5 to E16.5 in slice E1S1. We identified 2000 HVGs per slide using the Scanpy `highly_variable_genes` function.

The engineered mouse stem-cell-based embryo model data set comprises time-course scRNA-seq profiles tracking the lineage differentiation of control mouse gastruloids, derived solely from embryonic stem cells, and different ‘aggregoids’ (NACL, RACL, and XAL), derived by complementing embryonic cells with extraembryonic cells [24], from 48 h to 120 h at 24 h intervals. The data are quality controlled, integrated and annotated according to Smirnova et al. [24]. 2000 HVGs were identified across all samples using the Scanpy `highly_variable_genes` function.

## Results

### Mapping mouse development across time

To evaluate the robustness of scDIV against choices in computational workflows and variation in cell-type assignments, we compared the performance of both Hill numbers *D*_H_(*p*) and LC diversity *D*_LC_(*p*; *Z*) across multiple cell-type assignments of a single-cell time-lapse data set of mouse development [23] (Figure 2). We tested three distinct annotations representing different resolutions, software, and computational pipelines: (1) a coarse reference assignment consisting of 26 major cell clusters, (2) a fine-grained reference subclustering comprising 190 cell types, both derived via a Scanpy-based workflow from Qiu et al. [23], and (3) an independent, *de novo* coarse assignment of 40 clusters generated using Seurat in this study to simulate cross-laboratory and cross-platform pipeline variation. See Figure 2A–C.

**Figure 2.**
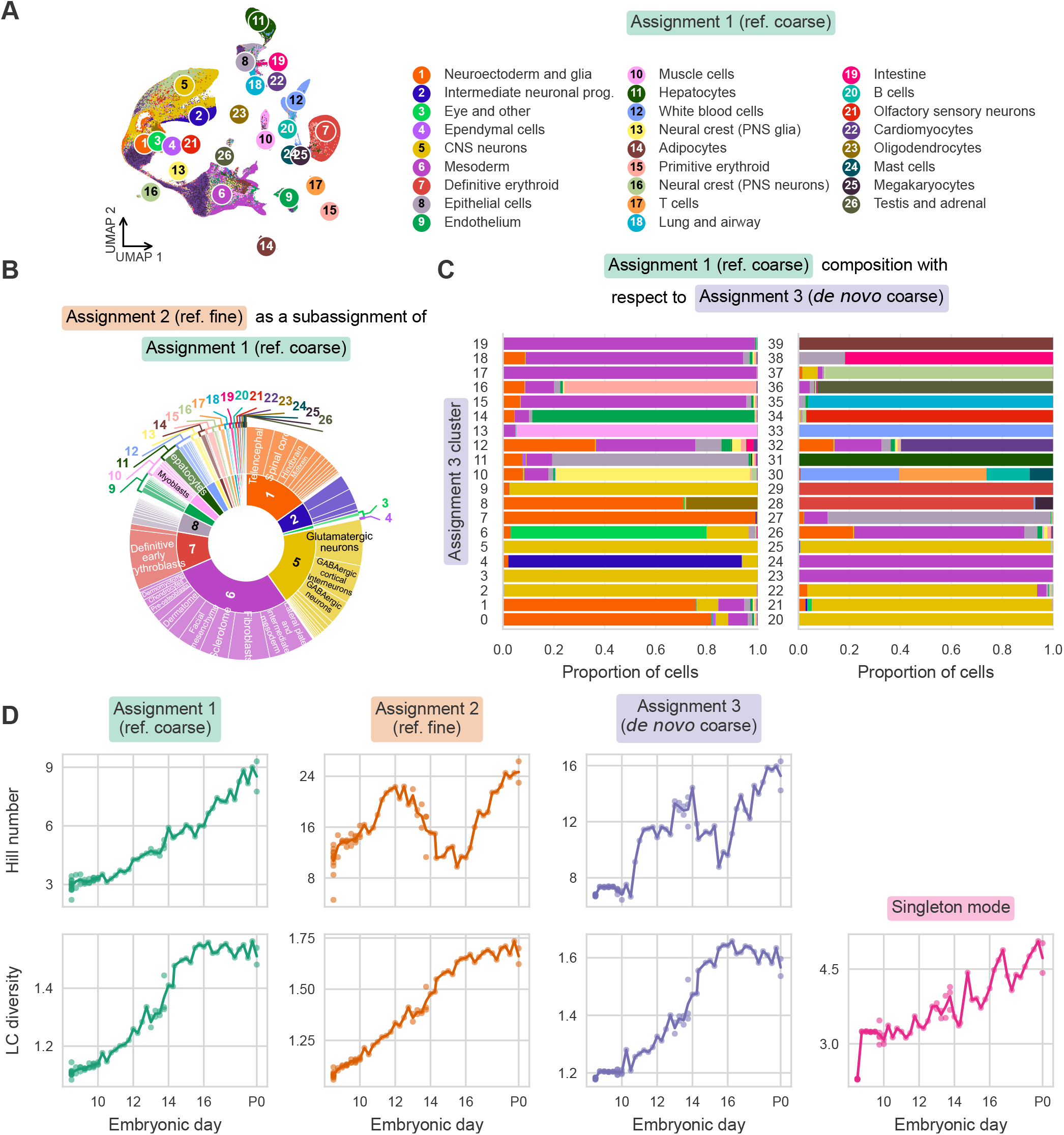
scDIV is robust against cell-type assignments. (A) Two-dimensional uniform manifold approximation and projection (UMAP) visualization of the mouse development data set. Color corresponds to Assignment 1 (coarse reference). Prog., progenitor. (B) Hierarchical lineage chart mapping the sub-assignment relationship between Assignment 1 (inner ring) and Assignment 2 (fine reference; outer ring). (C) Proportional bar chart mapping the relationship between Assignment 1 cell types (colored bars) relative to Assignment 3 clusters (*de novo* coarse; y-axis numbering). (D) Diversity trajectories across embryonic development (E8 to P0). Top row displays Hill numbers; bottom row displays LC diversity. Columns from left to right represent trajectories calculated using Assignment 1, Assignment 2, Assignment 3, and Singleton mode (LC diversity only). Solid lines pass through the mean diversity value at each timepoint, and individual points represent one mouse embryo.

We observed that the Hill number *D*_H_ fluctuates considerably depending on the choice of assignment (Figure 2D, top row). In contrast, LC diversity is robust to shifts in cell-type resolution. The LC diversity trajectories of the coarse reference (Assignment 1) and its highly partitioned subclustering (Assignment 2) demonstrated stability, maintaining an almost identical trend despite a seven-fold increase in the number of defined cell types. Furthermore, LC diversity proved to be highly reproducible across analysis software, demonstrating consistent trajectory trends between our Seurat-based *de novo* workflow (Assignment 3) and the Scanpy-based references.

Moving on to a biological interpretation, all three cell-type-based LC diversity scores successfully capture a rapid diversification of cell states immediately following gastrulation (E8.5). To further analyze this macro-level diversification, we calculated LC diversity in singleton mode, which bypasses predefined cell-type groupings entirely. Interestingly, while all cell-type-dependent trajectories eventually plateaued toward the end of embryonic development (after E16), the singleton-mode diversity continued to rise steadily in late development (Figure 2D, bottom right). This divergence suggests that, while the emergence of major canonical cell types stabilizes in late gestation, ongoing differentiation and maturation programs continue to diversify the transcriptional states of cells within established cell populations, as reflected by the continued increase in singleton-mode diversity.

As a further test of our robustness claims, we also compared LC diversity with the Hill number on the outputs of four different analysis pipelines on the same PBMC data set as computed by Rich et al. [6]. We found that LC diversity remains remarkably robust despite the differences in both cell type clustering and HVG set between the four pipelines. See Supplementary Figure S1.

### Breaking down local and global diversity *in situ*

Moving beyond single-cell time-series data, spatial transcriptomics allows us to map diversity dynamics *in situ* and examine where diversification processes occur across the physical architecture of the embryo. To investigate how cellular diversity is spatially organized across the developing mouse embryo, we applied our spatial diversity framework to the MOSTA Stereo-seq data set to calculate both local and global diversity (Figure 3).

**Figure 3.**
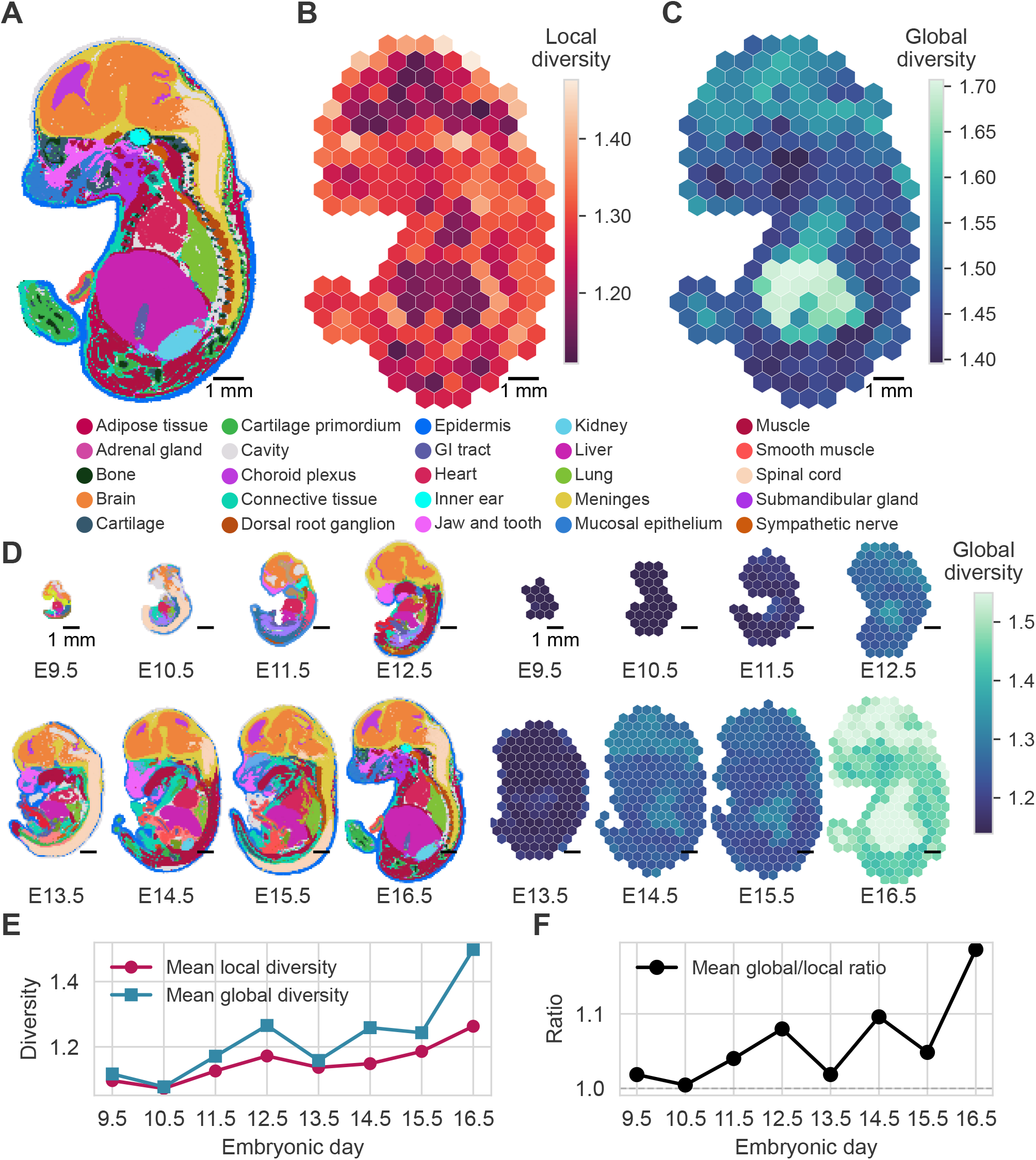
Spatial transcriptomics diversity dynamics across mouse organogenesis. (A) Spatial bin50 annotation of the E16.5 E1S1 mouse embryo section. Bins are colored according to their cell-type annotations (legend below). (B) Spatial distribution of local diversity calculated per region across the E16.5 mouse embryo section. (C) Spatial distribution of global diversity scores calculated per region across the E16.5 mouse embryo section. (D) Spatiotemporal atlas of mouse development and global diversity dynamics. Left: Time-series sagittal sections of mouse embryos across E9.5 to E16.5. Bins are colored by their cell-type annotations. Right: Corresponding spatial heat maps showing global diversity across the same embryonic stages (E9.5–E16.5). (E) Line plot tracking the mean local and global diversity as a function of the embryonic day. (F) Line plot tracking the mean global/local diversity ratio as a function of the embryonic day. Scale bars: 1 mm.

Computing the local and global diversity across individual regions of an E16.5 mouse section reveals that structural interfaces, where multiple cell lineages physically meet and mix, exhibit the highest local diversity. Conversely, large, expanding internal organs show much lower local diversity. An example is the developing liver, which displays a highly uniform, homogeneous cellular population. (Figure 3A, B)

While the liver is internally uniform, it exhibits a high global diversity score (Figure 3C). This combination of low local diversity and high global diversity indicates that the liver is composed of a specialized, highly distinct cell composition that is rarely found elsewhere in the embryo. This finding aligns with the macro-developmental lineage reported by Chen et al. [21]. In their characterization of anatomical relationships across developmental time, the liver stands out as an early-separating lineage branch from the endoderm germ layer. LC diversity successfully captures this phylogenetic divergence.

To understand how regional diversity emerges over developmental time, we examined the progression of global diversity across the full embryonic time series (Figure 3D). In early timepoints (E9.5–E11.5), the global diversity maps are nearly uniform and low in magnitude. This uniformity indicates that although early embryonic tissues are actively organizing, they have not yet diverged into specialized, compositionally distinct organs. By E12.5, the first signatures of regional specialization become apparent. Notably, developing internal organs, particularly the liver, begin to diverge transcriptomically from the surrounding tissue, marking the functional onset of organogenesis. This structural divergence becomes more visible at E16.5, where distinct regions of high global diversity emerge across the embryo.

Because our analysis considers only a single spatial section per embryonic timepoint, the data naturally reflects variations introduced by the specific anatomical plane of the cut, the proportion of cells captured, and individual biological variance between the mice. For instance, we observe a noticeable dip in both local and global diversity metrics at E13.5 (Figure 3E). Metadata from the original study suggests that this variation may be driven by lower sequencing depth compared to the adjacent timepoints as well as a more dominant section of the spinal cord captured.

Finally, the ratio of global to local diversity increases over time (Figure 3F). If development simply meant that every tissue generically became ‘more diverse’ over time, both local and global diversity would scale at roughly the same rate, keeping the ratio flat at 1.0. An increasing ratio demonstrates that the gain in global diversity is also to a large extent driven by spatial differentiation.

### Quantifying diversity in mouse embryo models

Stem-cell-based embryo models (SCBEMs) are powerful experimental systems to dissect interactions between the plethora of cell types that emerge during embryogenesis [25, 26]. Recent work leveraged the modularity of SCBEMs to test the effect of extra-embryonic endoderm on cellular composition and patterning by co-aggregation of embryonic and extra-embryonic cells (termed ‘aggregoids’) [24]. Compared to the control condition (gastruloids solely derived from embryonic stem cells), the aggregoids – termed NACL/RACL/XAL dependent on the type of extra-embryonic endoderm-like cells added – were reported to display a higher cell-type diversity based on traditional graph-based clustering and subsequent cell-type annotation [24]. We computed cell-type LC diversity, singleton LC diversity as well as Hill numbers to quantify the shifts in transcriptomic diversity of engineered aggregoids (Figure 4).

**Figure 4.**
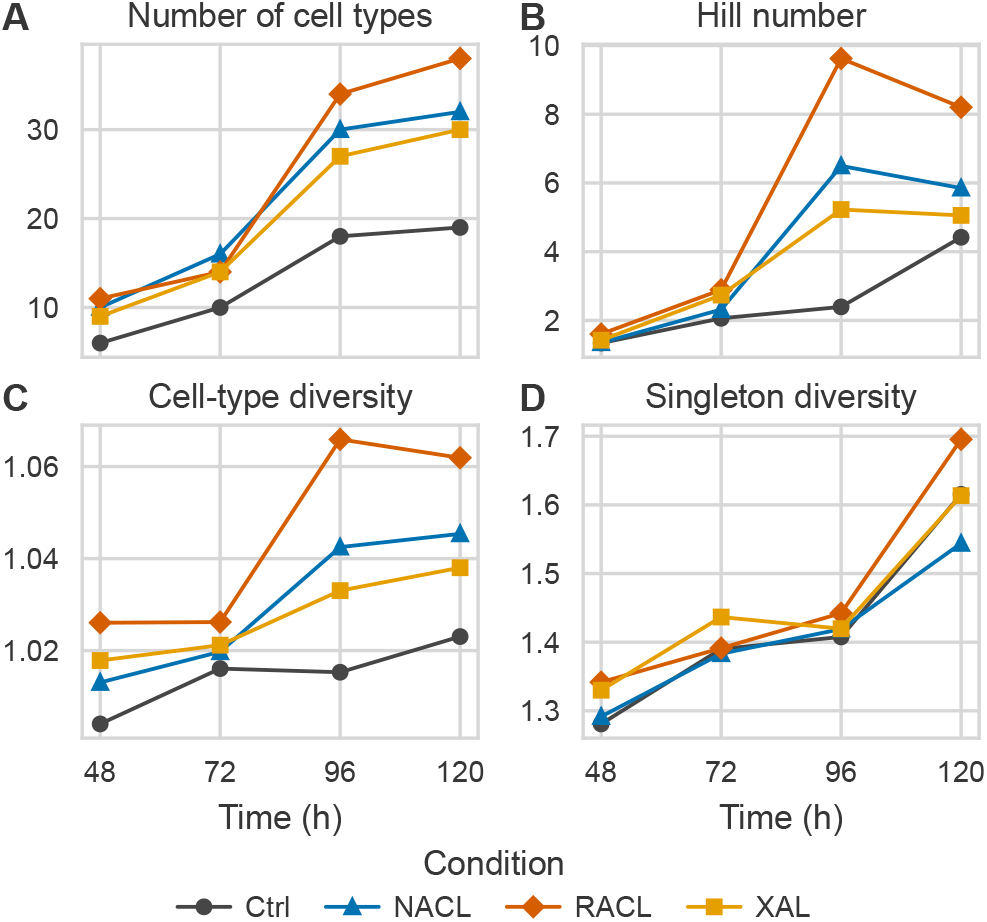
scDIV measures of engineered embryo models. (A–D) Line plots of diversity metrics across four engineered embryo conditions (Ctrl, NACL, RACL, and XAL) measured at 48 h, 72 h, 96 h and 120 h. (A) Total number of distinct cell types identified per sample. (B) Hill numbers. (C) cell-type LC diversity. (D) Singleton LC diversity.

Across all timepoints, LC diversity values remained within a relatively low overall range, reflecting the less distinct nature of the emerging cell types and the fact that these early-stage cells have not yet reached terminal differentiation states. At the 48-hour mark, both the cell-type-based and singleton LC diversity metrics, alongside traditional Hill numbers, effectively detected the initial introduction of the added extra-embryonic cells. By 120 hours, the cell-type diversity across the modified aggregoid conditions captured an expansion in overall composition driven by this extra-embryonic addition. Notably, however, singleton diversity uniquely revealed that the NACL condition deviated from this trend, demonstrating a lower singleton diversity relative to the control. This sensitivity is further highlighted between 90 and 120 hours, where all aggregoid conditions exhibited decoupled dynamics: while the Hill numbers decreased and cell-type LC diversity remained nearly constant, singleton diversity steadily increased.

## Discussion

Taken together, our results establish scDIV, a similarity-sensitive LC diversity framework, as a robust approach for quantifying cellular heterogeneity across single-cell and spatial transcriptomics datasets. At a higher level, this methodology provides a reliable mathematical metric for tracking both temporal lineage diversification and multi-scale spatial specialization within complex tissues. By decoupling the measurement of transcriptomic diversity from the subjective boundaries of manual cell-type annotation, scDIV opens up a reproducible path for benchmarking engineered biological models, mapping continuous cellular maturation, and comparing massive organismal atlases.

While the primary focus of this work has been on transcriptomic data, the mathematical formulation of LC diversity is fundamentally generalizable. Because the framework depends only on abundance distributions and a well-defined pairwise similarity matrix, it can be naturally extended to other single-cell data modalities. For instance, appropriate similarity measures could be defined for chromatin accessibility profiles (scATAC-seq), surface protein abundance (CITE-seq), or spatial morphometric features. Furthermore, this inherent flexibility makes LC diversity uniquely well-suited for multi-omics and multi-modal data sets, where a joint similarity matrix could capture multifaceted cellular identities simultaneously without requiring discrete, multi-modal clustering.

From a methodological standpoint, an important practical consideration is the effect of feature selection on the metric. We observe that LC diversity exhibits sensitivity when comparing pipelines that utilize all genes against those that restrict the feature space to highly variable genes (HVGs). This sensitivity arises because the inclusion of either housekeeping genes or highly sparse genes alters the underlying cosine similarity calculations between cells. However, within a reasonable and standard operational range of HVG choices, LC diversity remains highly stable. To ensure consistent and biologically meaningful diversity estimates, we recommend that users follow standard, data-set-specific preprocessing pipelines to select HVGs prior to running the computation.

### Limitations of the study

The computed LC diversity score should not be directly interpreted as an optimal ‘effective number’ of cell types to target in upstream clustering algorithms. In ecology, an LC score of *N* represents the complexity of *N* entirely distinct, equally abundant species. In transcriptomics, however, no universal consensus exists for defining a cell type, and ideal annotation boundaries vary across data sets and analytical goals. Because LC diversity is entirely agnostic to these biological conventions, its value serves as an abstract metric of continuous transcriptomic heterogeneity rather than a guide for cell-type partitioning choices.

## Resource availability

### Lead contact

Requests for further information and resources should be directed to and will be fulfilled by the lead contact, Heather A Harrington.

### Materials availability

This study did not generate new materials.

### Data and code availability

Source data statement: this paper analyzes existing data. These accession numbers for the data sets are

- The mouse development single-cell atlas is publicly available on the NCBI Gene Expression Omnibus (GEO) under accession number GSE228590.
- The mouse organogenesis spatial transcriptomics atlas is publicly available on the CNGB Nucleotide Sequence Archive under accession code CNP0001543.
- The scRNA-seq data set of engineered stem-cell-based embryo models has been deposited in the NCBI Gene Expression Omnibus (GEO) and will be made publicly accessible upon publication of the corresponding study.

Code statement: The Python software package `scdiv` is publicly available on GitHub at https://github.com/katherine-benjamin/scdiv. All original experimental code will be made available upon publication.

#### Acknowledgments

We thank Natalia Smirnova and Stefan Krauss (Oslo University Hospital and Hybrid Technology Hub Centre of Excellence, Oslo, Norway) for early access to the aggregoid data. JVV and HAH are funded by the Max Planck Gesellschaft. Work in JVV lab is funded by a European Innovation Council (EIC) Pathfinder grant under the Horizon Europe Research and Innovation Program (Horizon-EIC-2021-PathfinderChallenges-01 101071203, SUMO). HAH gratefully acknowledges funding from the Royal Society RGF*\*EA*\*201074, UF150238 and EPSRC EP/Y028872/1 and EP/Z531224/1. HAH and KB are grateful for the support provided by the UK Centre for Topological Data Analysis EPSRC grant EP/R018472/1. KB was supported by the JT Hamilton and EPSRC scholarship. LJ is a member of the International Max Planck Research School hosted at the MPI-CBG/CSBD. For the purpose of Open Access, the authors have applied a CC BY public copyright licence to any author accepted manuscript (AAM) version arising from this submission.

## Declaration of generative AI and AI-assisted technologies

During the preparation of this work, the authors used Claude Opus 4.8 in order to develop features and tests for the `scdiv` software package and to write the Python scripts which generated the figures in the manuscript. After using this tool or service, the authors reviewed and edited the content as needed and take full responsibility for the content of the publication.

## References

1. H. Kitano. Biological robustness. Nat. Rev. Genet., 5(11): 826–837, Nov. 2004. 10.1038/nrg1471.

2. J. Stelling, U. Sauer, Z. Szallasi, F. J. Doyle, and J. Doyle. Robustness of cellular functions. Cell, 118(6):675–685, Sept. 2004. 10.1016/j.cell.2004.09.008.

3. R. Elmentaite, C. Domínguez Conde, L. Yang, and S. A. Teichmann. Single-cell atlases: shared and tissue-specific cell types across human organs. Nat. Rev. Genet., 23(7):395–410, July. 2022. 10.1038/s41576-022-00449-w.

4. N. Schaum, J. Karkanias, N. F. Neff, A. P. May, S. R. Quake, T. Wyss-Coray, S. Darmanis, J. Batson, O. Botvinnik, M. B. Chen et al. Single-cell transcriptomics of 20 mouse organs creates a Tabula Muris. Nature, 562(7727):367–372, Oct. 2018. 10.1038/s41586-018-0590-4.

5. X. Han, Z. Zhou, L. Fei, H. Sun, R. Wang, Y. Chen, H. Chen, J. Wang, H. Tang, W. Ge et al. Construction of a human cell landscape at single-cell level. Nature, 581(7808):303–309, May. 2020. 10.1038/s41586-020-2157-4.

6. J. M. Rich, L. Moses, P. H. Einarsson, K. Jackson, L. Luebbert, A. S. Booeshaghi, S. Antonsson, D. K. Sullivan, N. Bray, P. Melsted et al. The impact of package selection and versioning on single-cell RNA-seq analysis. Cell Syst., 17(4):101560, Apr. 2026. 10.1016/j.cels.2026.101560.

7. T. Leinster and C. A. Cobbold. Measuring diversity: the importance of species similarity. Ecology, 93(3):477–489, Mar. 2012. 10.1890/10-2402.1.

8. T. Leinster and M. W. Meckes. Maximizing diversity in biology and beyond. Entropy, 18(3):88, Mar. 2016. 10.3390/e18030088.

9. S. Mohammadi, J. Davila-Velderrain, and M. Kellis. A multiresolution framework to characterize single-cell state landscapes. Nat. Commun., 11(1):5399, Oct. 2020. 10.1038/s41467-020-18416-6.

10. J. C. van Wolfswinkel, D. E. Wagner, and P. W. Reddien. Single-cell analysis reveals functionally distinct classes within the planarian stem cell compartment. Cell Stem Cell, 15(3):326–339, Sept. 2024. 10.1016/j.stem.2014.06.007.

11. A. Heindl, C. Lan, D. N. Rodrigues, K. Koelble, and Y. Yuan. Similarity and diversity of the tumor microenvironment in multiple metastases: critical implications for overall and progression-free survival of high-grade serous ovarian cancer. Oncotarget, 7(44):71123–71135, 2016. 10.18632/oncotarget.12106.

12. A.-M. Tsakiroglou, S. Astley, M. Dave, M. Fergie, E. Harkness, A. Rosenberg, M. Sperrin, C. West, R. Byers, and K. Linton. Immune infiltrate diversity confers a good prognosis in follicular lymphoma. Cancer Immunol. Immunother., 70(12):3573–3585, 2021. 10.1007/s00262-021-02945-0.

13. C. R. S. Banerji, D. Miranda-Saavedra, S. Severini, M. Widschwendter, T. Enver, J. X. Zhou, and A. E. Teschendorff. Cellular network entropy as the energy potential in waddington’s differentiation landscape. Sci. Reports, 3, 2013. 10.1038/srep03039.

14. A. E. Teschendorff and T. Enver. Single-cell entropy for accurate estimation of differentiation potency from a cell’s transcriptome. Nat. Commun., 8(1):15599, June. 2017. 10.1038/ncomms15599.

15. A. Richard, L. Boullu, U. Herbach, A. Bonnafoux, V. Morin, E. Vallin, A. Guillemin, N. P. Gao, R. Gunawan, J. Cosette et al. Single-cell-based analysis highlights a surge in cell-to-cell molecular variability preceding irreversible commitment in a differentiation process. PLOS Biol., 14(12):e1002585, Dec. 2016. 10.1371/journal.pbio.1002585.

16. T. T. Karagiannis, S. Monti, and P. Sebastiani. Cell Type Diversity Statistic: An Entropy-Based Metric to Compare Overall Cell Type Composition Across Samples. Front. Genet., 13:855076, Apr. 2022. 10.3389/fgene.2022.855076.

17. T. Leinster. Entropy and Diversity: the Axiomatic Approach. Cambridge University Press, Cambridge, United Kingdom, 2021.

18. M. O. Hill. Diversity and Evenness: A Unifying Notation and Its Consequences. Ecology, 54(2):427–432, Mar. 1973. 10.2307/1934352.

19. M. T. Seweryn, M. Pietrzak, and Q. Ma. Application of information theoretical approaches to assess diversity and similarity in single-cell transcriptomics. Comput. Struct. Biotechnol. J., 18:1830–1837, 2020. 10.1016/j.csbj.2020.05.005.

20. R. Reeve, T. Leinster, C. A. Cobbold, J. Thompson, N. Brummitt, S. N. Mitchell, and L. Matthews. How to partition diversity. arXiv:1404.6520 [q-bio]. 10.48550/arXiv.1404.6520, Dec. 2016.

21. A. Chen, S. Liao, M. Cheng, K. Ma, L. Wu, Y. Lai, X. Qiu, J. Yang, J. Xu, S. Hao et al. Spatiotemporal transcriptomic atlas of mouse organogenesis using DNA nanoball-patterned arrays. Cell, 185(10):1777–1792.e21, May. 2022. 10.1016/j.cell.2022.04.003.

22. R. R. Stickels, E. Murray, P. Kumar, J. Li, J. L. Marshall, D. J. Di Bella, P. Arlotta, E. Z. Macosko, and F. Chen. Highly sensitive spatial transcriptomics at near-cellular resolution with Slide-seqV2. Nat. Biotechnol., 39(3): 313–319, 2021. 10.1038/s41587-020-0739-1.

23. C. Qiu, B. K. Martin, I. C. Welsh, R. M. Daza, T.-M. Le, X. Huang, E. K. Nichols, M. L. Taylor, O. Fulton, D. R. O’Day et al. A single-cell time-lapse of mouse prenatal development from gastrula to birth. Nature, 626:1084–1093, Feb. 2024. 10.1038/s41586-024-07069-w.

24. N. P. Smirnova, S. V. Ponomartsev, T. M. L. Ali, M. Lycke, B. K. Chung, J. Øgaard, E. Melum, T. Combriat, J. V. Veenvliet, and S. K. Deiml. Modular engineering of embryonic-extraembryonic interactions generates advanced gastruloid morphotypes. bioRxiv:2025.11.09.687163. 10.1101/2025.11.09.687163, June. 2026.

25. J. Fu, A. Warmflash, and M. P. Lutolf. Stem-cell-based embryo models for fundamental research and translation. Nat. Mater., 20(2):132–144, Feb. 2021. 10.1038/s41563-020-00829-9.

26. M. Bao, J. Cornwall-Scoones, and M. Zernicka-Goetz. Stem-cell-based human and mouse embryo models. Curr. Opin. Genet. & Dev., 76:101970, Oct. 2022. 10.1016/j.gde.2022.101970.

