## Supplemental text and figures for "Diversity in transcriptomics without cell types"

S1.4 A note on computing singleton-mode diversity with cosine similarity 4

**Supplementary Figures 7**

### S1 Similarity-sensitive diversity in detail

This note contains further details on the computation of similarity-sensitive diversity measures. For a full exposition on diversity measures we refer the reader to Leinster (2021).

#### S1.1 Similarity-sensitive diversity of order $q$

In the main text (Materials and Methods: Similarity-sensitive diversity) we introduced the LC diversity measure. In fact, this measure is a single instantiation of a broader family of measures known as the LC diversities *parametrized by an order* $q \in [0, \infty]$. The `scdiv` package exposes the parameter $q$ to the user, and we explain here how it operates.

We begin by introducing the notion of a power mean of order $q$.

**Definition S1.** For a distribution $p \in [0, 1]^N$ and a vector $x \in [0, \infty)^N$, with $x_i > 0$ whenever $i \in \text{supp}(p)$, the *power mean of order $q$ of $x$, weighted by $p$*, is defined piecewise. For $q \notin \{-\infty, 0, \infty\}$ it is given by

$$M_q(p; x) = \left( \sum_{i \in \text{supp}(p)} p_i x_i^q \right)^{1/q} . \tag{S1}$$

Additionally, we have

$$M_{-\infty}(p; x) = \min_{i \in \text{supp}(p)} x_i, \tag{S2}$$

$$M_0(p; x) = \prod_{i \in \text{supp}(p)} x_i^{p_i}, \tag{S3}$$

$$M_{\infty}(p; x) = \max_{i \in \text{supp}(p)} x_i. \tag{S4}$$

Now we give the definition of LC diversity in full generality.

**Definition S2.** Let $(\mathcal{S}, p, Z)$ be a community. For $q \in [0, \infty]$ its *LC diversity of order* $q$ is

$$D^q_{\text{LC}}(p; Z) \coloneqq M_{1-q}(p; 1/(Zp)) \tag{S5}$$

where $(1/(Zp))_i = 1/(Zp)_i$.

Note that we have

$$D^2_{\text{LC}}(p; Z) = M_{-1}(p; 1/(Zp)) = \left( \sum_{i \in \text{supp}(p)} p_i (Zp)_i \right)^{-1} \tag{S6}$$

and this is precisely the LC diversity as defined in the main text. So in the main text we consider only diversity of order 2.

One can think of the order $q$ as controlling the *insensitivity* of the diversity measure to rare members of the community. At the extreme, when $q = \infty$ we care only about the least-unique member. So by setting $q = 2$ in the experiments in this paper we have been assigning a relatively high importance to rare cell types.

#### S1.2 Stability

The following two properties of LC diversity give us the robustness under subclustering that we make use of in the main text. The first property says that LC diversity is indifferent to trivial subdivisions of cell types into identical subtypes (Leinster, 2021, Lemma 6.2.6).

**Proposition S3.** *Let $(\mathcal{S}, p, Z)$ be a community with members $\mathcal{S} = [S_1, \ldots, S_{N+1}]$. Write $(\mathcal{S}', p', Z')$ for the community with members $\mathcal{S}' = [S_1, \ldots, S_{N-1}, S'_N]$ obtained by merging $S_N$ and $S_{N+1}$ into a single member $S'_N$, so that $p'$ agrees with $p$ on $S_1, \ldots, S_{N-1}$ and assigns abundance $p_N + p_{N+1}$ to $S'_N$. If $S_N$ and $S_{N+1}$ are indistinguishable within the community, i.e. $Z_{Ni} = Z_{(N+1)i}$ for all $i$, then $D^q_{\text{LC}}(p'; Z') = D^q_{\text{LC}}(p; Z)$ for all $q \in [0, \infty]$.*

The second property says that LC diversity is stable with respect to small changes to the similarity matrix (Leinster, 2021, Lemma 6.2.4(ii)).

**Proposition S4.** *Fix $q \in [0, \infty]$ and a distribution $p$. Then $D^q_{\text{LC}}(p; Z)$ is continuous in $Z$.*

Combining Propositions S3 and S4 gives the robustness to subtyping we use extensively in the main text. In particular, moving from a 'supertype' clustering to a 'subtype' clustering involves first splitting the supertype into identical subtypes, which does not change diversity at all, and then distorting the similarity matrix to match the new subtypes. If the subtypes are reasonably similar to the supertype, then this distortion will only have a mild effect on diversity thanks to the continuity of diversity in the similarity matrix $Z$.

##### S1.3 Partitioned diversity

Suppose we have a community that is subdivided into some collection of disjoint subcommunities. There are two natural ways in which we might consider a subcommunity to be diverse: either it consists of a diverse population of members as a community in itself, or it contributes strongly to the diversity of the total community by containing many unique members relative to the global population.

Reeve et al. (2016) formalized these ideas in the context of LC diversity by introducing the notion of *partitioned diversity*. Consider a metacommunity $(\mathcal{S}, p, Z)$, which is divided into $N$ subcommunities. Let $P_{ij}$ represent the abundance of member $i$ in subcommunity $j$ relative to the total metacommunity, thus defining an $S \times N$ abundance matrix $P$ where the column vector $P_{\cdot j} = (P_{1j}, \ldots, P_{Sj})$ represents the raw relative abundances within subcommunity $j$. The total weight or relative size of subcommunity $j$ is denoted by $w_j = \sum_i P_{ij}$, and its isolated, normalized type distribution is given by $\overline{P}_{\cdot j} = P_{\cdot j}/w_j$.

We now introduce the *local diversity*[1] $\overline{\alpha}_j^q(Z)$, which quantifies the heterogeneity of subcommunity $j$ viewed in isolation. It represents the effective number of members within the subcommunity after controlling for its size.

**Definition S5.** Let $(\mathcal{S}, p, Z)$ be a metacommunity partitioned into $N$ subcommunities with relative abundance matrix $P$. For a subcommunity $j \in \{1, \ldots, N\}$, the *local diversity* of order $q$ is defined as:

$$\overline{\alpha}_j^q(Z) := M_{1-q}\left(\overline{P}_{\cdot j}, 1/Z\overline{P}_{\cdot j}\right) \tag{S7}$$

where $(Z\overline{P}_{\cdot j})_i = \sum_{i'} Z_{ii'}\overline{P}_{i'j}$ represents the ordinariness of type $i$ within the isolated subcommunity $j$.

In the main text, we consider the local diversity of order $q = 2$,

$$\overline{\alpha}_j^2(Z) = M_{-1}\left(\overline{P}_{\cdot j}, 1/Z\overline{P}_{\cdot j}\right) = \left(\sum_{i:\overline{P}_{ij}>0} \overline{P}_{ij}(Z\overline{P}_{\cdot j})_i\right)^{-1}. \tag{S8}$$

Another notion of subcommunity diversity is the *global diversity*[2], $\gamma_j^q(Z)$, which measures the contribution per individual within subcommunity $j$ toward the

[1] Which Reeve *et al.* call the *normalized-alpha diversity.*

[2] Which Reeve *et al.* call the *gamma diversity.*

total diversity of the metacommunity. It is calculated by weighting the global uniqueness of the members by the subcommunity's relative distribution:

**Definition S6.** Let $(\mathcal{S}, p, Z)$ be a metacommunity partitioned into $N$ subcommunities with relative abundance matrix $P$. For a subcommunity $j \in \{1, \ldots, N\}$, the *global diversity* of order $q$ is defined as:

$$\gamma_j^q(Z) := M_{1-q}\left(\overline{P}_{\cdot j}, 1/Zp\right) \tag{S9}$$

where $(Zp)_i = \sum_{i'} Z_{ii'} p_{i'}$ represents the ordinariness of type $i$ across the entire global metacommunity.

Again, in the main text, we consider the global diversity of order $q = 2$,

$$\gamma_j^2(Z) = M_{-1}\left(\overline{P}_{\cdot j}, 1/Zp\right) = \left(\sum_{i \in \mathrm{supp}(p)} \overline{P}_{ij}(Zp)_i\right)^{-1}. \tag{S10}$$

In the context of single-cell transcriptomics, the entire tissue sample serves as the metacommunity, whereas individual subcommunities may be specific cell types, germ layers, or organs. When spatial coordinates are available, as in spatial transcriptomics, subcommunities can be taken to be localized spatial domains, such as anatomical layers or grid-based spatial bins.

##### S1.4 A note on computing singleton-mode diversity with cosine similarity

Computing diversity involves evaluating the matrix product $Zp$ where $Z \in M_N([0,1])$ is our similarity matrix and $p \in [0,1]^N$ is our relative abundance distribution. In singleton-mode, $N$ is the number of cells and it is therefore undesirable to instantiate the dense $N \times N$ matrix $Z$. Luckily, the fact that we are using cosine similarity allows us to avoid instantiating the entire matrix.

Recall that the cosine similarity for a gene expression matrix $X$ is given by

$$Z := \overline{X}^T \overline{X} \tag{S11}$$

where $\overline{X}_j = X_j / \|X_j\|$. The product structure means that we can compute the product $Zp$ by

$$Zp = (\overline{X}^T \overline{X})p = \overline{X}^T(\overline{X}p) \tag{S12}$$

and avoid computing the $N \times N$ similarity matrix by instead first computing the $N \times 1$ column vector $\overline{X}p$. In fact, in singleton mode we have $p = (1/N, \ldots, 1/N)^T$ so that $(\overline{X}p)_i = \frac{1}{N}\sum_{j=1}^n \overline{X}_{ij}$.

### References

Tom Leinster. *Entropy and Diversity: the Axiomatic Approach.* Cambridge University Press, Cambridge, United Kingdom, 2021.

Richard Reeve, Tom Leinster, Christina A. Cobbold, Jill Thompson, Neil Brummitt, Sonia N. Mitchell, and Louise Matthews. How to partition diversity, December 2016. URL `http://arxiv.org/abs/1404.6520`.

Joseph M. Rich, Lambda Moses, Pétur Helgi Einarsson, Kayla Jackson, Laura Luebbert, A. Sina Booeshaghi, Sindri Antonsson, Delaney K. Sullivan, Nicolas Bray, Páll Melsted, and Lior Pachter. The impact of package selection and versioning on single-cell RNA-seq analysis. *Cell Systems*, 17(4):101560, April 2026. doi: 10.1016/j.cels.2026.101560.

### Supplementary Figures

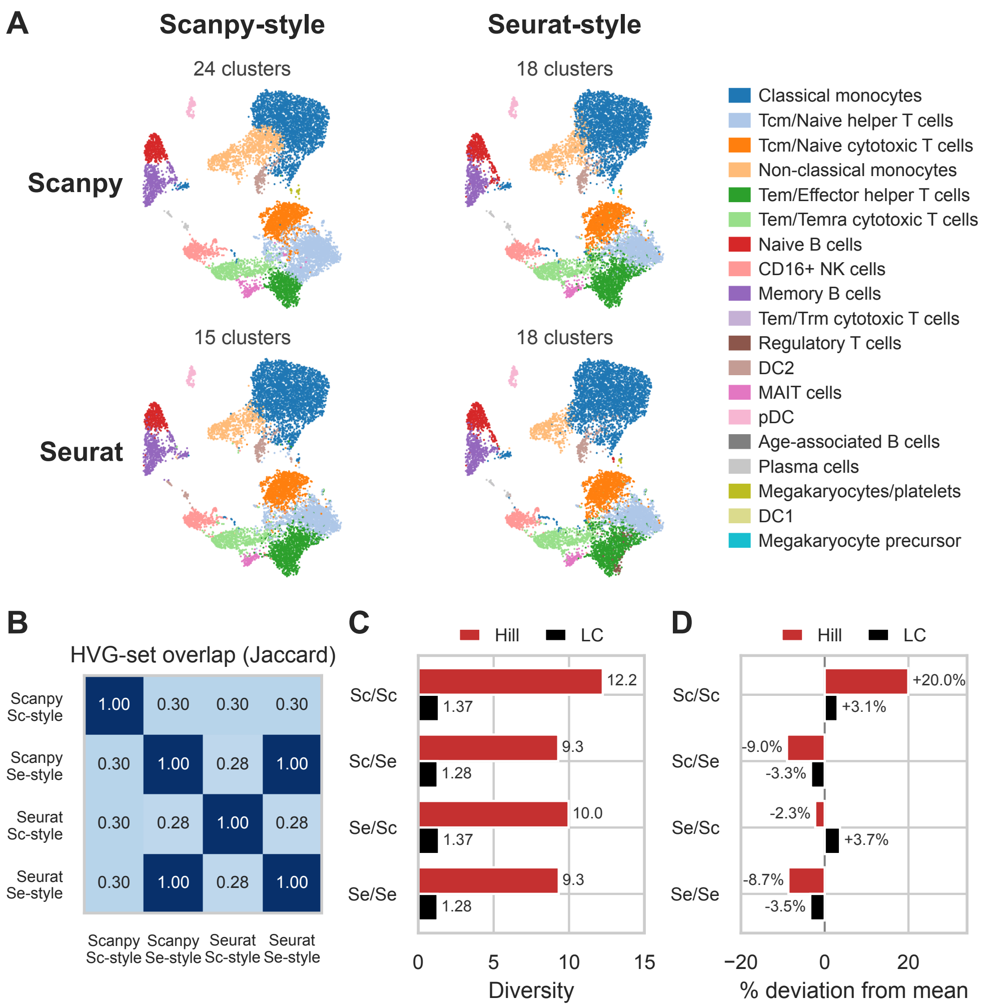

Figure S1: LC diversity of four PBMC clusterings from Rich et al. (2026). Each pipeline fixes a tool and a parameter set from either Seurat (Se) or Scanpy (Sc). (A) A UMAP projection of the data set colored by each clustering. Clusters are colored by their majority cell type in a shared CellTypist cell-type labeling. (B) Jaccard overlap of the four 2000-gene HVG sets from each pipeline. (C) Hill number and LC diversity of each clustering and HVG set. (D) Deviation from the mean of each diversity measure. The LC diversity remains robust to relabeling.
